# TBX3 acts as tissue-specific component of the Wnt/β-catenin enhanceosome

**DOI:** 10.1101/2020.04.22.053561

**Authors:** Dario Zimmerli, Costanza Borrelli, Amaia Jauregi-Miguel, Simon Söderholm, Salome Brütsch, Nikolaos Doumpas, Jan Reichmuth, Fabienne Murphy-Seiler, Michel Aguet, Konrad Basler, Andreas E. Moor, Claudio Cantù

**Affiliations:** Department of Molecular Life Sciences, University of Zurich, Zürich, Switzerland, CH-8057; Institute of Molecular Cancer Research, University of Zurich, Zürich, Switzerland, CH-8057; Wallenberg Centre for Molecular Medicine, Linköping University; Department of Biomedical and Clinical Sciences, Faculty of Health Science, SE-581 83 Linköping, Sweden; Swiss Institute for Experimental Cancer Research (ISREC), Ecole Polytechnique Fédérale de Lausanne (EPFL), School of Life Sciences, CH-1015 Lausanne, Switzerland; Division of Molecular Pathology, The Netherlands Cancer Institute, Amsterdam, The Netherlands

## Abstract

BCL9 and PYGO are β-catenin cofactors that enhance the transcription of Wnt target genes. They have been proposed as therapeutic targets to diminish Wnt signalling output in intestinal malignancies. Here we find that, in colorectal cancer cells and in developing mouse forelimbs, BCL9 proteins sustain the action of β-catenin in a largely PYGO-independent manner. Our genetic analyses implied that BCL9 necessitates other interaction partners in mediating its transcriptional output. We identified the transcription factor TBX3 as a candidate tissue-specific member of the β-catenin transcriptional complex. In developing forelimbs, TBX3 and BCL9 co-occupy a large number of Wnt-responsive regulatory elements, genome-wide. Moreover, mutations in *Bcl9* affect the expression of TBX3 targets in vivo, and modulation of TBX3 abundance impacts on Wnt target genes transcription in a β-catenin- and TCF/LEF-dependent manner. Finally, TBX3 overexpression exacerbates the metastatic potential of Wnt-dependent human colorectal cancer cells. Our work implicates TBX3 as a new, context-dependent component of the Wnt/β-catenin-dependent enhanceosome.

## Introduction

The Wnt pathway is an evolutionarily conserved cell signalling cascade that acts as major driving force of several developmental processes, as well as for the maintenance of the stem cell populations within adult tissues (Nusse and Clevers, 2017). Deregulation of this signalling pathway results in a spectrum of consequences, ranging from lethal developmental abnormalities to several forms of aggressive cancer (Nusse and Clevers, 2017). Most prominently, colorectal cancer (CRC) is initiated by genetic mutations that constitutively activate Wnt signalling (Kahn, 2014).

Secreted WNT ligands trigger an intracellular biochemical cascade in the receiving cells that culminates in the calibrated expression of target genes (Mosimann et al., 2009). This transcriptional response is orchestrated by nuclear β-catenin, that acts as a “scaffold” to buttress a host of co-factors to cis-regulatory elements occupied by the TCF/LEF transcription factors (Valenta et al., 2012). Among the co-factors, the two paralogs BCL9 and BCL9L (referred to as BCL9/9L) and PYGO1/2 proteins reside within the so-called Wnt enhanceosome, and their concerted action is required to efficiently activate Wnt-target gene expression (Kramps et al., 2002; Parker et al., 2002; van Tienen et al., 2017) (Figure 1A). During vertebrate development, their requirement in the β-catenin-mediated transcription appears to be context-dependent (Cantù et al., 2018; Li et al., 2007), and they also have evolved β-catenin-independent functions (Cantù et al., 2017, 2014). Curiously however, BCL9 and PYGO always seem to act as a “duet” (Kennedy et al., 2010).

**Figure 1:**
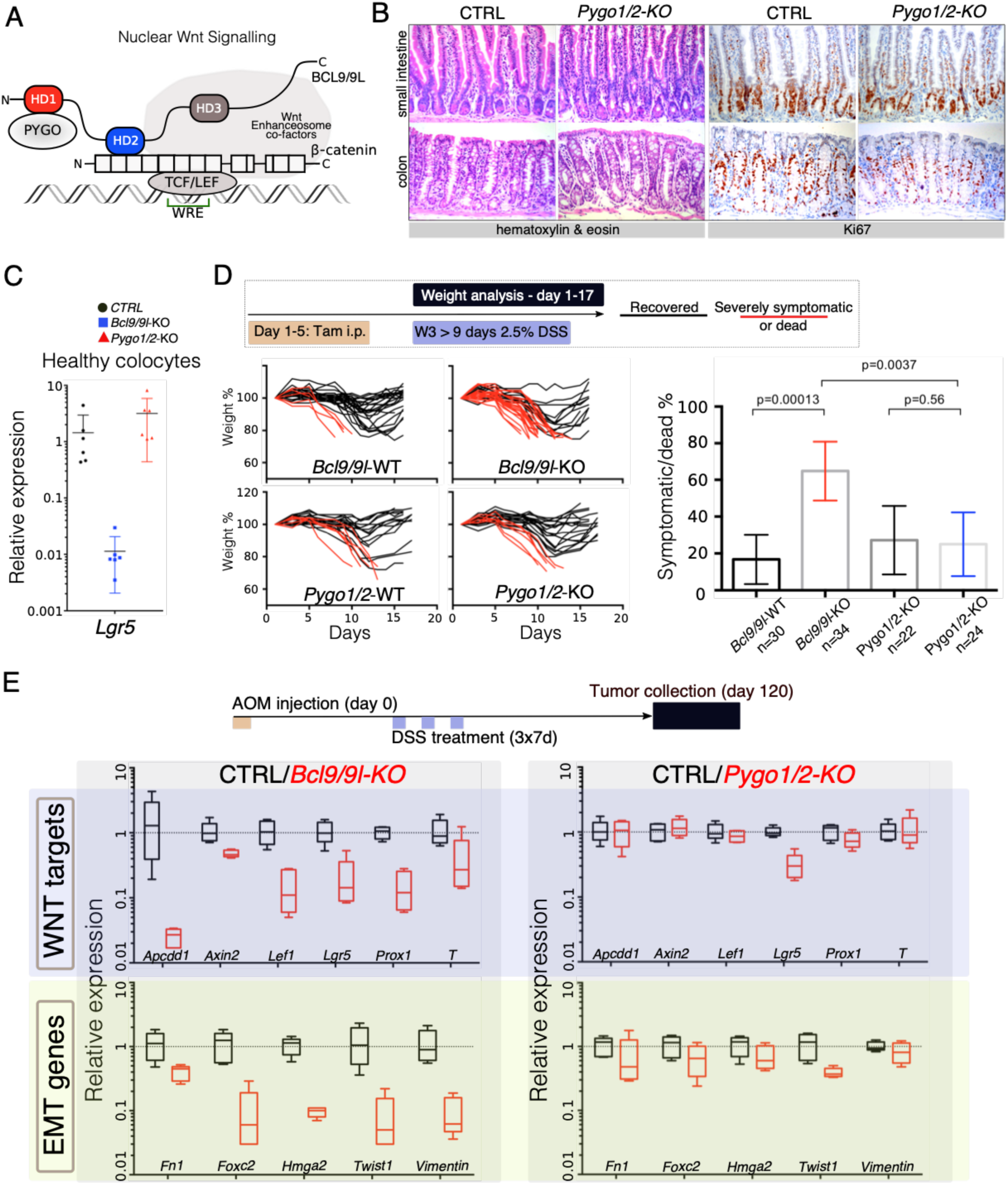
The intestinal epithelium-specific recombination of *Pygo1/2* does not recapitulate the effects of deleting *Bcl9/9l*. (**A**) Schematic representation of the Wnt enhanceosome, with emphasis on the so-called “chain of adaptors” components, - catenin, BCL9/9L and PYGO1/2; WRE, Wnt Responsive Element. The homology domains 1-3 (HD1-3) of BCL9/9L are shown. (**B**) Epithelial-specific *Pygo1/2* deletion (via *vil-Cre-ER*^*t2*^; *Pygo1/2*-KO) does not lead to any obvious histological or functional defect, neither in the small intestine nor in the colon. Intestinal architecture was normal, crypts were present and contained proliferating positive cells as seen by hematoxylin and eosin staining (left panels), and count of the proliferative compartment via Ki67 (right panels), respectively. (**C**) Quantitative RT-PCR detecting *Lgr5* mRNA extracted from colonic epithelium of control (black), *Bcl9/9l* (blue) or *Pygo1/2* (red) conditional mutants (KO). (**D**) 6-8-week-old male mice were treated with five tamoxifen (Tam) injections (i.p., 1mg/day) for five consecutive days. 10 days later mice were treated with 2.5% dextran sodium sulfate (DSS) ad libitum in the drinking water for 9 days. While 17% of control mice (n=30) were severely affected or died due to the DSS treatment (red lines), 65% of conditional *Bcl9/9l*-KO (n=34) mice performed poorly in this test. Deletion of *Bcl9/9l* increased statistically significantly the death rate after DSS treatment (p-value = 0.00013 in Fisher’s Exact Test). No difference between *Pygo1/2*-KO and control mice could be measured: 27% of control mice (n=22) and 25% of *Pygo1/2*-KO (n=24) were affected upon DSS treatment (p-value = 0.5626 in Fisher’s Exact Test). (**E**) 6-8-week-old female mice were exposed to a single dose of the carcinogenic agent azoxymethane (AOM), followed by 7 days of DSS administration in the drinking water. This regimen results in the emergence of dysplastic adenomas that are collected for RNA extraction and analysis of the indicated targets via RT-PCR: Wnt target genes and genes expressed during epithelial-to-mesenchymal transition (EMT), associated with cancer metastasis.

Importantly, BCL9/9L and PYGO proteins were found to significantly contribute to the malignant traits typical of Wnt-induced CRCs (Deka et al., 2010; Gay et al., 2019; Jiang et al., 2020; Mani et al., 2009; Mieszczanek et al., 2019; Moor et al., 2015; Talla and Brembeck, 2016). These observations provided impetus to consider the BCL9/PYGO axis as relevant “targetable” unit in CRC (Lyou et al., 2017; Mieszczanek et al., 2019; Talla and Brembeck, 2016; Zimmerli et al., 2017).

However, here we noticed an apparent divergence between the roles of BCL9/9L and PYGO proteins. We found that genetic abrogation of *Bcl9/9l* in mouse CRC cells results in broader consequences than *Pygo1/2* deletion, suggesting that BCL9 function does not entirely depend on PYGO1/2. Among the putative β-catenin/BCL9 interactors we identified the developmental transcription factor TBX3. Intriguingly, we show that also during forelimb development, BCL9/9L possess a PYGO-independent role. In this in vivo context, TBX3 co-occupies β-catenin/BCL9 target loci genome-wide, and mutations in *Bcl9/9l* affect the expression of TBX3 targets. Finally, TBX3 modulates the expression of Wnt target genes in a β-catenin- and TCF/LEF-dependent manner, and increases the metastatic potential of human CRC cells when overexpressed. We conclude that TBX3 can assist the Wnt/β-catenin mediated transcription in selected developmental contexts, and that this partnership could be aberrantly reactivated in some forms of Wnt-driven CRCs.

## Results and discussion

We induced intestinal epithelium-specific recombination of *Pygo1/2* loxP alleles (*Pygo1/2*-KO), that efficiently deleted these genes in the whole epithelium, including the stem cells compartment (Figure supplement 1A and 1B). Consistently with recent reports (Mieszczanek et al., 2019; Talla and Brembeck, 2016), and similarly to deletion of *Bcl9/9l* (Deka et al., 2010; Mani et al., 2009; Moor et al., 2015), *Pygo1/2*-KO displayed not overt phenotypic defects (Figure 1B and Figure supplement 1C). We were surprised in noticing that the expression of *Lgr5*, the most important intestinal stem cell marker and Wnt target gene (Barker et al., 2007), was heavily downregulated upon loss of *Bcl9/9l* but unaffected in *Pygo1/2*-KO (Figure 1C). To address the functionality of the stem cell compartment in these two conditions, we subjected both *Bcl9/9l* and *Pygo1/2* compound mutants (KO) to a model of intestinal regeneration by DSS treatment (Kim et al., 2012) (Figure 1D). While *Bcl9/9l*-KO mice showed a defect in regeneration after insult (Deka et al., 2010), *Pygo1/2*-KO proved indifferent when compared to control littermates (Figure 1D). While we cannot exclude that PYGO1/2 have subtle transcriptional contribution, our results highlight that the BCL9/9L function in the intestinal epithelium homeostasis and regeneration does not entirely depend on PYGO1/2. This was surprising, since BCL9/9L proteins were thought to act as mere “bridge” proteins that tethered PYGO to the β-catenin transcriptional complex (Figure 1A) (Fiedler et al., 2015; Mosimann et al., 2009). Both BCL9 and PYGO proteins have been implicated in colorectal carcinogenesis (Gay et al., 2019; Jiang et al., 2020; Mieszczanek et al., 2019; Talla and Brembeck, 2016). We tested if the consequence of the deletion of *Bcl9/9l* and *Pygo1/2* genes was also different in the context of carcinogenesis. Specifically, we looked at the contribution to gene expression in chemically-induced AOM/DSS colorectal tumors (Figure 1E). As previously observed, *Bcl9/9l*-KO tumors exhibit a massive decrease in Wnt target gene expression, EMT and stemness traits (Deka et al., 2010; Moor et al., 2015), which was not observed in *Pygo1/2*-KO tumors (Figure 1E, Figure supplement 2). This phenotypic difference is consistent with a recent study in which *Bcl9/9l* but not *Pygo1/2* loss reduced the activation of Wnt target genes induced by *APC* loss-of-function (Mieszczanek et al., 2019); we interpret this as an independent validation of our observation. All these experiments open up the question of how BCL9/9L imposes its function independently of PYGO.

Surprisingly, the intestine-specific deletion of the homology domain 1 (HD1) of BCL9/9L (Figure 2A), that was previously annotated to interact only with PYGO1/2 (Cantù et al., 2014; Kramps et al., 2002), induced i) beneficial phenotypic changes that are not observed upon deletion of *Pygo1/2* (Figure 2B) and ii) a strong downregulation of Wnt target, EMT and stemness genes (Figure 2C, figure supplement 2C). The discrepancy between the gene expression changes induced by recombining *Pygo1/2* or deleting the HD1 domain of *Bcl9/9l* implies that currently unknown proteins assist BCL9/9L function. We set out to identify new candidate BCL9 partners that might be responsible for the different phenotypes. To this aim, we performed a pull-down of tumor proteins expressing either a full-length or a HD1-deleted variant of BCL9, followed by mass spectrometry (Figure 2D). Among the proteins differentially pulled down by control but not by mutant BCL9 we detected TBX3 (Figure 2D and 2E) and selected it for further validation. TBX3 appeared as the most interesting candidate, since the malformations induced by *Bcl9/9l* loss-of-function are strikingly similar to those induced by *Tbx3* loss (Frank et al., 2013). Indeed, the in vivo deletion of the HD1 domain (in *Bcl9/9l*- ΔHD1 embryos) leads to severe forelimb malformations, while *Pygo1/2-*KO embryonic forelimbs are unaffected (Figure 2F)(also see Schwab et al., 2007). Limb development, thus, represents another context where BCL9/9L appear to act independently of PYGO.

**Figure 2:**
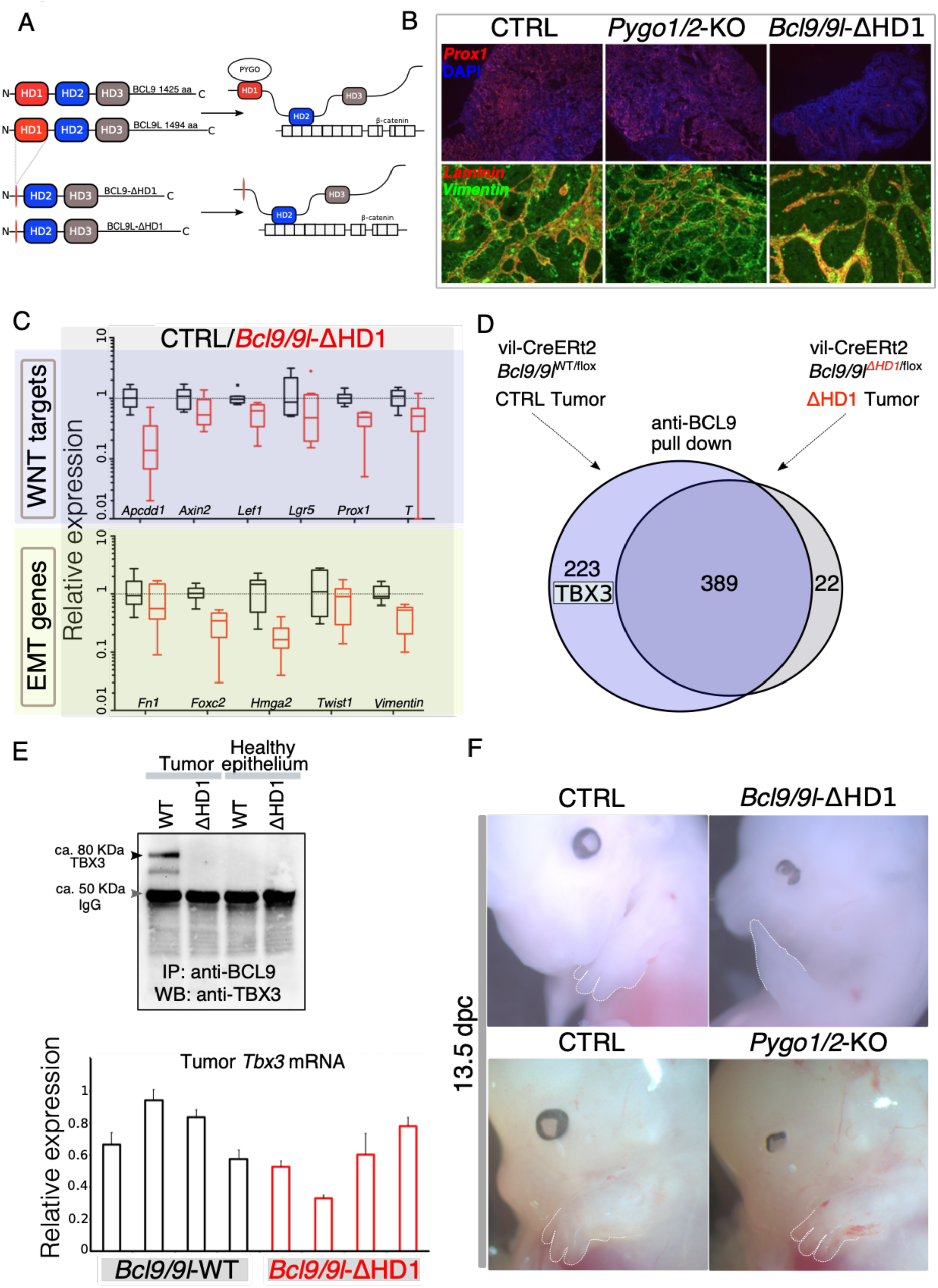
Identification of TBX3 as a putative BCL9 cofactor. (**A**) The deletion of the HD1 (PYGO-interacting) domain of BCL9 and BCL9L induces a variation in the “chain of adaptors” implying loss of PYGO association with the Wnt enhanceosome (Cantù et al., 2014). (**B**) Immunofluorescence staining of tumors collected from control or conditional Pygo1/2-KO and *Bcl9/9l-*KO mice. Prox1 (red) and DAPI (blue) are shown in in the top panels; Vimentin (green) and Laminin (red) in the bottom panels. (**C**) Quantitative RT-PCR of select groups of targets (compare it with the same analysis of *Pygo1/2*-KO in figure 1E) of RNA extracted from control or *Bcl9/9l*-ΔHD1 tumors. (**D**) Experimental outline of the tumor proteins pull-down and mass-spectrometry. TBX3 was identified among the proteins potentially interacting with BCL9 but not with BCL9-ΔHD1. (**E**) The protein IP analyzed by mass spectrometry were in parallel subjected to SDS page electrophoresis and probed with an anti-TBX3 antibody (upper panel). The expression of *Tbx3* in control compared to Bcl9/9l-ΔHD1 tumors (N=4) was evaluated via qRT-PCR (bottom panel), to exclude that differential pull-down was due to lost expression in mutant tumors. (**F**) 13.5 dpc Bcl9/9l-ΔHD1 embryos display forelimb malformations and absence of digits (emphasized by dashed white lines) – a characteristic *Tbx3*-mutant phenotype (upper panels). The limb defect is absent in Pygo1/2-KO embryos (bottom panels) underscoring that BCL9/9L act, in this context, independently of PYGO1/2.

We confirmed cytological vicinity between transfected tagged versions of BCL9 and TBX3 by proximity ligation assay (PLA) (Figure supplement 3A). However, overexpression-based in vitro co-immunoprecipitation experiments could not detect any stable interaction between these two proteins, suggesting absence of direct binding or a significantly lower affinity than that between BCL9 and PYGO (Figure supplement 3B). Hence, we aimed at testing the functional association between TBX3 and BCL9 in a more relevant in vivo context. To this aim, we collected ca. 500 forelimbs from 10.5 dpc wild-type mouse embryos and subjected the crosslinked chromatin to immunoprecipitation using antibodies against BCL9 (Salazar et al., 2019) or TBX3, followed by deep-sequencing of the purified DNA (ChIP-seq, Figure 3A). By using stringent statistical parameters, and filtering with Irreproducible Discovery Rate (IDR), we extracted a list of high confidence BCL9 and TBX3 peaks (Figure 3B and 3C). Surprisingly, we discovered that BCL9 co-occupies a large fraction (ca. 2/3rd) of the TBX3-bound regions (Figure 3D). Suggestive of a role for TBX3 within the Wnt-dependent transcriptional apparatus, motif analysis of the co-occupied loci identified statistical prevalence for TCF/LEF and Homeobox transcription factors consensus sequences, but not that for any TBX transcription factor (Figure E). This suggests that TBX3 interacts with the DNA in these locations via affinity to the Wnt enhanceosome rather than via direct contact with DNA. Accordingly, TBX-specific motifs were detected within the group of TBX3 exclusive peaks (which are not co-occupied by BCL9, Figure supplement 4). Notably, TBX3 and BCL9 co-occupancy was detected at virtually all previously described Wnt-responsive-elements (WRE) within known Wnt target genes (Figure 3F).

**Figure 3:**
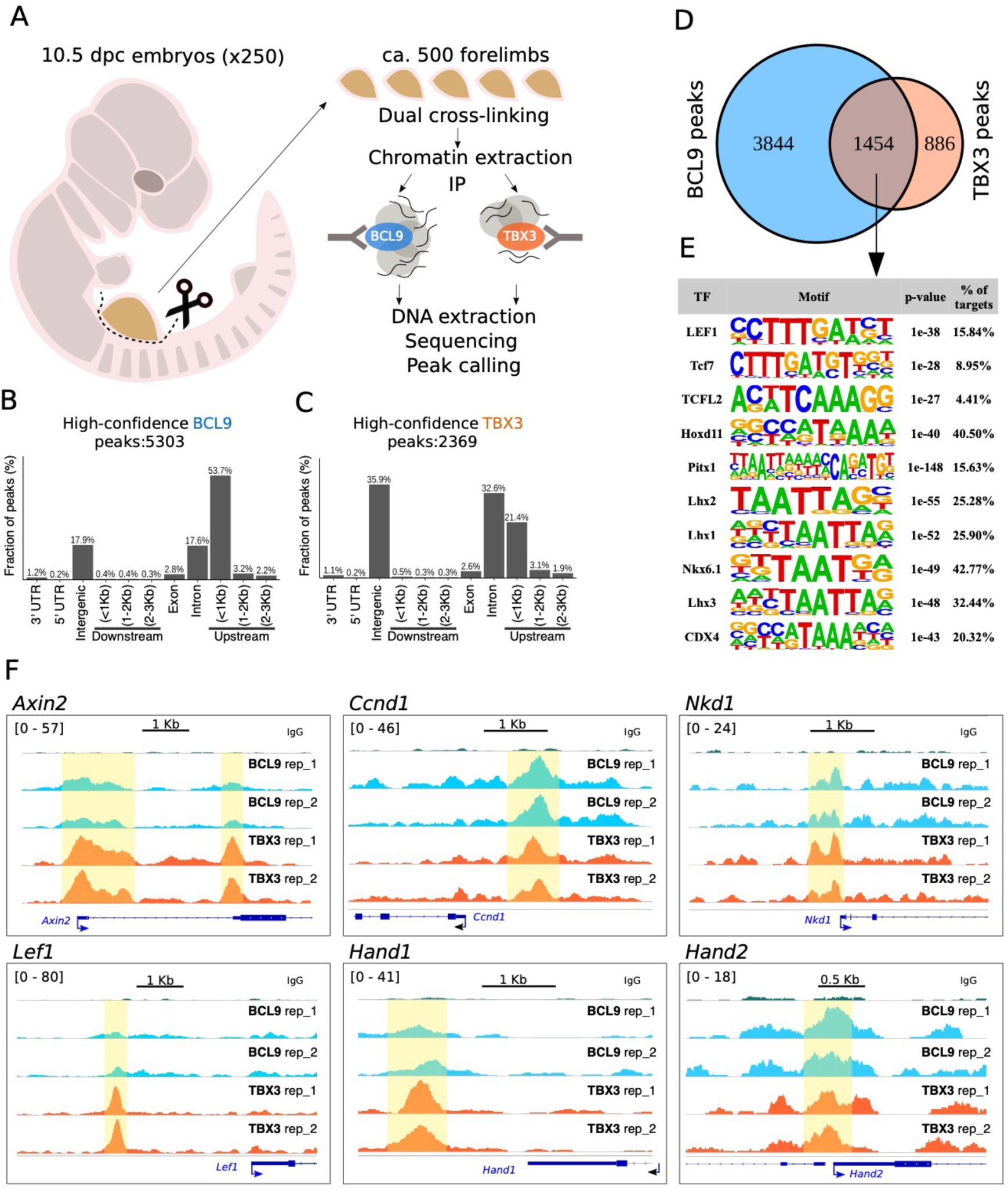
TBX3 and BCL9 co-occupy Wnt Responsive Elements (WRE) in vivo. (**A**) Artistic representation of the ChIP-seq experimental outline. (**B-C**) Bar-plots showing the genomic distribution of high-confidence BCL9 peaks (B, 5303 total) and TBX3 peaks (C, 2369 total). (**D**) Overlap of the high-confidence peak groups between BCL9 and TBX3. (**E**) Selected result entries from motif analysis performed on the BCL9-TBX3 overlapping high-confidence peaks. Significant enrichment was found for TCF/LEF and Homeobox motifs. No TBX consensus sequence was detected in this analysis. (**F**) Select genomic tracks demonstrating co-occupancy of BCL9 and TBX3 within the Wnt Responsive Element (WRE) of known Wnt-target genes (*Axin2*, *Ccnd1*, *Nkd1* and *Lef1*) and genes important in limb morphogenesis (*Hand1* and *Hand2*). The scale of peak enrichment is indicated in the top-left corner of each group of tracks. In light blue the BCL9 (Salazar et al., 2019), in orange the TBX3 replicates, and in green the control track (IgG). Genomic tracks are adapted for this figure upon visualization with IGV Integrative Genomic Viewer (https://igv.org/).

So far, we have presented genetic evidence that BCL9 proteins require additional co-factors, and that TBX3 associates with the β-catenin/BCL9 bound regions on the genome. We reasoned that our hypothesis - in which BCL9 functionally tethers TBX3 to the β-catenin transcriptional complex - raises several testable predictions that will be addressed below. First, if TBX3 is physically tethered by BCL9 on its targets, mutations in *Bcl9/9l* should influence the expression of genes associated with TBX3 peaks. Second, our model implies that TBX3 could impact on Wnt target genes expression and its activity should be dependent on the presence of both β-catenin and TCF/LEF. Finally, as for BCL9, TBX3 should be capable of enhancing the metastatic potential of colorectal cancer cells.

To test our first prediction, that is whether mutations in *Bcl9/9l* would influence genes associated with TBX3 peaks, we set out to mutate *Bcl9/9l* interaction domains, while leaving TBX3 protein unaffected. We generated mouse embryos bearing genetic abrogation of both the HD2 (β-catenin-interacting) and HD1 (PYGO-interacting) motifs (*Bcl9/9l*-Δ1/Δ2; Figure 1A) Cantù et al., 2018); note that this genetic configuration induces forelimbs malformations that are PYGO-independent, as previously shown (Figure 2F). We collected forelimbs from control and *Bcl9/9l*-Δ1/Δ2 mutant embryos at 10.5, and measured gene expression via RNA-seq (Figure 4A). Strikingly, we found a significant enrichment (Hypergeometric test, p=1.4e-6) of TBX3 targets among the genes differentially expressed in *Bcl9/9l*-Δ1/Δ2 mutants (Figure 4B). The enrichment was particularly significant when considering down-regulated genes in *Bcl9/9l*-Δ1/Δ2 mutants, indicating that the BCL9-TBX3 partnership sustains transcriptional activation (Figure 4B). Of note, the design of our experiment directly implicates that these TBX3 transcriptional targets are also β-catenin-dependent. The overlap list includes several regulators of limb development, such as *Meis2* (Capdevila et al., 1999), *Irx3* (Li et al., 2014) and *Eya2* (Grifone et al., 2007) (Figure 4B, heat map on the right).

**Figure 4:**
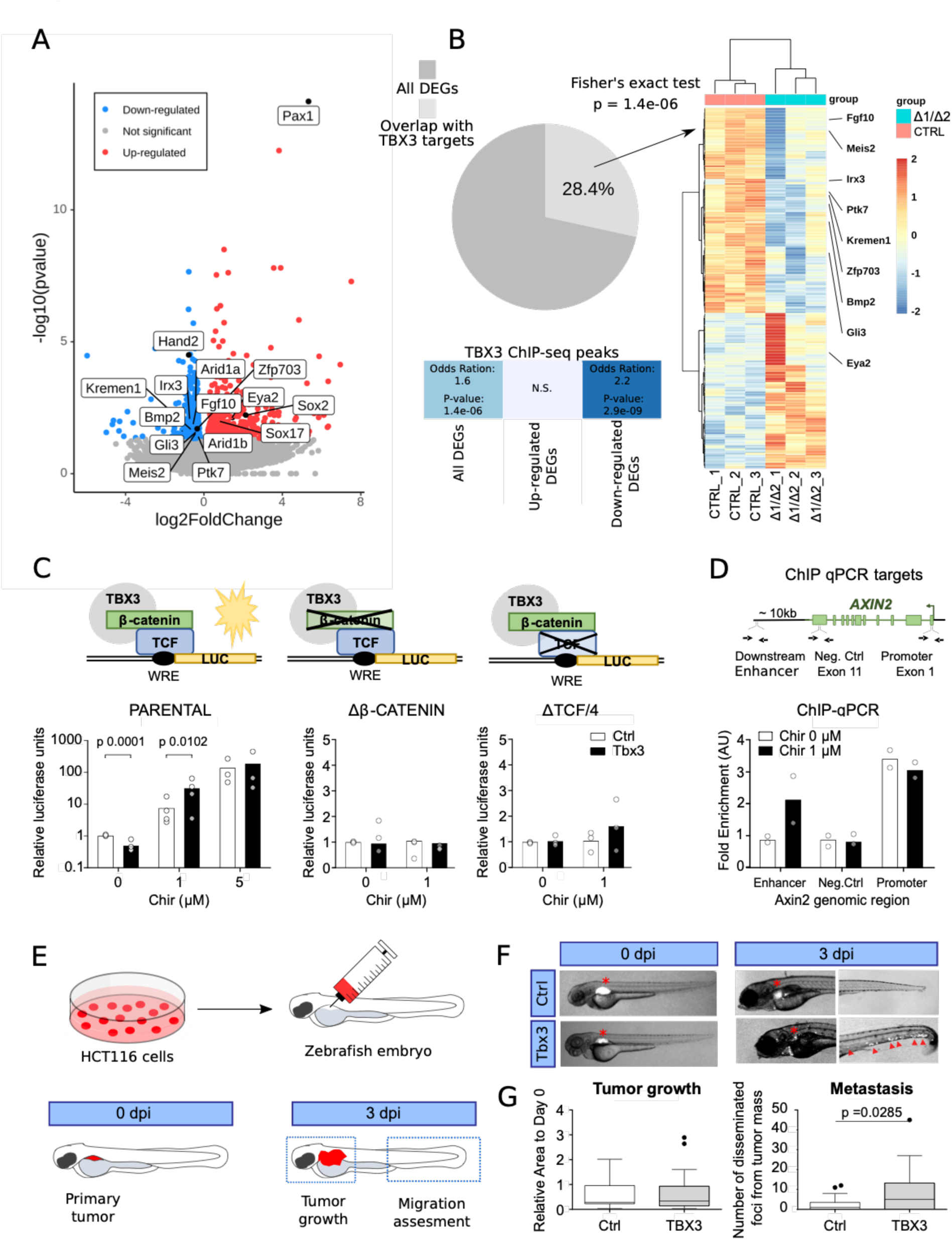
TBX3 controls the expression of Wnt target genes. (**A**) Volcano plot displays all the differentially expressed genes (DEGs) in developing forelimbs, upon mutation of *Bcl9/9l* (*Bcl9/9l-Δ*1/Δ2 vs CTRL). DEGs were a total of 1143 (p < 0.05), with 606 up-regulated and 537 down-regulated. (**B**) A significant portion (28.4%) of DEGs exhibited overlap with TBX3 ChIP-seq peaks. The overlap with TBX3 ChIP-seq peaks appeared statistically significant, in particular, when the down-regulated genes were considered. (**D**) Hierarchical clustering of samples (3 CTRL versus 3 *Bcl9/9l-Δ*1/Δ2) based on genes overlapping between DEGs and genes annotated for TBX3 ChIP-seq peaks (normalized RNA-seq read counts, Ward’s clustering method, Euclidian distance). Annotation added for genes associated by Gene Ontology to Wnt signaling (*Fgf10*, *Ptk7*, *Kremen1*, *Zfp703*, *Bmp2* and *Gli3*) and genes known as regulators of limb development (*Meis2*, *Irx3* and *Eya2*). (**C**) β-catenin/TCF luciferase reporter STF assay in parental, β-catenin knockout and TCF knockout HEK293T cells. Cells were treated with the indicated concentration of CHIR or DMSO, overnight. Expression of TBX3 (black bars) compared to control (empty vector, white bars) showed that TBX3 acts as an activation switch for Wnt/TCF pathway. Only significant p-values (p<0.05) are indicated. (**D**) ChIP followed by qPCR in HEK293T cells treated with DMSO (“WNT-OFF”) or CHIR (“WNT-ON”). Schematic representation of the *AXIN2* locus indicates the position of the primers used (black arrows) to test binding of TBX3. Enrichment was identified on *AXIN2* promoter and the downstream enhancer. Data are normalized to immunoprecipitation with a control IgG antibody and presented as the mean ± standard deviation of technical duplicate. (**E**) Schematic diagram of the human CRC zebrafish xenografts model. Parental and TBX3-overexpressing HCT116 colorectal tumor cells were harvested and labeled with DiI dye (red). The stained cells were injected into the perivitelline space of 3-day old zebrafish embryos. Zebrafish were visualized with fluorescent microscopy at 0-day post injection (dpi) and 3 dpi, and primary tumor cell invasion and metastasis were counted. (**F**) Representative images of HCT116 tumor invasion and dissemination at 0 and 3 dpi in zebrafish xenografts, for both control and TBX3 overexpressing cells. The red asterisks indicate the position of the primary tumor. Red arrowheads point at clusters of disseminating/metastatic cells. (**G**) Quantification of primary tumor growth and metastasis after HCT116 xenograft. Data are presented as mean ± standard deviation. Only significant p-values (p<0.05) are displayed.

We then addressed our second prediction implying a potential role of TBX3 in the transcription of Wnt target genes. We overexpressed it into HEK293T cells and monitored the activation status of Wnt signalling using the transcriptional reporter SuperTopFlash (STF). Consistent with its role as repressor, TBX3 led to a moderate but significant transcriptional downregulation that was, importantly, specific to the STF but not the control reporter plasmid (Figure 4C). Upon Wnt signalling activation achieved via GSK3 inhibition, TBX3-overexpressing cells exhibited a markedly increased reporter activity when compared to control cells, in particular at non-saturating pathway stimulating conditions (Figure 4C). Importantly, TBX3 proved transcriptionally incompetent on the STF if the cells carried mutations in *TCF*/*LEF* or *CTNNB1* (encoding for β-catenin; Doumpas et al., 2018), strongly supporting the notion of its cell-autonomous involvement in the activation of canonical Wnt target gene transcription (Figure 4C, central and right panels, respectively). Endogenous Wnt targets showed a similar expression behaviour to that of STF upon TBX3 overexpression (Figure supplement 5). Consistently, TBX3 was bound to *Axin2* promoter both in “OFF” and in “ON” conditions (Figure 4D).

Finally, we evaluated the effects of TBX3 overexpression (OE) on growth and metastatic potential of HCT116 human colorectal tumor cells – a representative model of CRC driven by activating mutations in *CTNNB1* (Mouradov et al., 2014) -, using a in vivo zebrafish xenograft model (Rouhi et al., 2010). Approximately 200-500 labelled control or TBX3-OE HCT116 cells were implanted in the perivitelline space of 72 hours post-fertilization (hpf) zebrafish embryos (Figure 4E). Three days after injection, TBX3-OE cells displayed a marked increase in number, both at the injection site and in the caudal hematopoietic plexus (Figure 4F and 4G), the main metastatic site for cells migrating from the perivitelline space (Rouhi et al., 2010). This indicated that TBX3 increases proliferation and migratory capability of human CRC cells bearing a constitutively active Wnt signalling.

Taken together, our experiments show that, in specific developmental and disease contexts, the transcription factor TBX3 can take active part in the direct regulation of Wnt target genes by functional interplay with the β-catenin/BCL9-dependent transcriptional complex. Our study suggests a new paradigm in which tissue-specific co-factors might be the key to understand the spectrum of possible transcriptional outputs observed downstream of Wnt/β-catenin signalling. Moreover, TBX3 has been linked to different cancer types (Willmer et al., 2017). Our observations suggest that TBX3, or its downstream effectors, could be considered as new relevant targets to dampen CRC progression.

## Materials and Methods

### Treatment of mice and histological analyses

#### Homeostasis

4-6-week-old male and female mice (*Bcl9f*^flox/flox^;*Bcl9l*^flox/flox^; vil-Cre-ERt2 and *Bcl9f*^flox/flox^;*Bcl9l*^flox/flox^ (no Cre) littermates; *Pygo1f*^flox/flox^;*Pygo2*^flox/flox^; vil-Cre-ERt2 and *Bcl9f*^flox/flox^;*Bcl9l*^flox/flox^ (no Cre)) were treated with five tamoxifen injections (i.p., 1mg/day, Sigma) for five consecutive days and the small intestine and colon were analyzed at different time points thereafter. Mouse experiments were performed in accordance with Swiss guidelines and approved by the Veterinarian Office of Vaud, Switzerland.

Induction of DSS Colitis: 6-8-week-old male mice were treated with five tamoxifen injections (i.p., 1mg/day) for five consecutive days. 10 days later 2.5% DSS (MG 36-50’000, MP Biomedicals, cat. no. 160110) was administered, ad libitum, in the drinking water for 9 days.

Induction of tumors: 6-8-week-old female mice were treated with five tamoxifen injections (i.p., 1mg/day) for five consecutive days. 10 days later they were injected i.p. with 44mg/kg body weight DMH 2HCl (N,N’ Dimethylhydrazine dihydrochloride). After another 7 days later 2% DSS was administered, ad libitum, in the drinking water for 7 days.

Mice were monitored clinically for rectal bleeding, prolapse and general signs of morbidity, including hunched posture, apathetic behavior and failure to groom.

The relative body weight (in %) was calculated as follows: 100 X weight at a certain day / weight at the first day of DSS treatment. Epithelial damage of DSS treated mice was defined as percentage of distal colon devoid of epithelium.

To determine proliferation rates, mice were injected i.p. with 100mg/kg BrdU (Sigma) 2hr prior to sacrifice. Small intestines and colons (divided into three equal segments to be named proximal, middle and distal colon) were dissected, flushed with cold PBS, cut open longitudinally and fixed in 4% paraformaldehyde for 2hr at RT and paraffin embedded. Sections (4µm) were cut and used for hematoxylin/eosin and alcian blue staining and for immunohistochemistry. The primary antibodies used were rabbit anti-Synptophysin (DAKO; 1:100), rabbit anti-Lysozyme (1:500; DAKO), mouse anti-Ki67 (1:100; Novocastra), mouse anti-BrdU (1:500; Sigma), anti-β-catenin (1:100; BD pharmigen), anti-active caspase 3 (1:100; Cell Signaling).

The peroxidase-conjugated secondary antibodies used were Mouse or Rabbit EnVision+ (Dako) or mouse anti-rat HRP (1:250: Biosource).

### Real-time PCR genotyping

To determine the deletion rate, the intestinal epithelium was separated from the underlining muscle. The intestine was dissected, flushed with PBS, cut open longitudinally and incubated in 3mM ethylenediamine tetraacetic acid (EDTA) and 0.05mM dithiothreitol (DTT) in PBS for 1.5h at RT on a rotor. The tubes were shaken vigorously, the muscle removed, and the epithelium centrifuged and used for genomic DNA extraction. SYBR green real-time PCR assays were performed on each sample analyzed. Genotyping primers are available upon request.

### Chromatin Immunoprecipitation

Forelimb buds were manually dissected from ca. 250 RjOrl:SWISS outbred 10.5 dpc mouse embryos. Chromatin immunoprecipitation was performed as previously described (Cantù et al., 2018). Briefly, the tissue was dissociated to a single cell suspension with collagenase (1 ug/ml in PBS) for 1 hr at 37° C, washed and crosslinked in 20 ml PBS for 40 min with the addition of 1.5 mM ethylene glycol-bis(succinimidyl succinate) (Thermo Scientific, Waltham, MA, USA), for protein-protein crosslinking (Salazar et al., 2019), and 1% formaldehyde for the last 20 min of incubation, to preserve DNA-protein interactions. The reaction was blocked with glycine and the cells were subsequently lysed in 1 ml HEPES buffer (0.3% SDS, 1% Triton-X 100, 0.15 M NaCl, 1 mM EDTA, 0.5 mM EGTA, 20 mM HEPES). Chromatin was sheared using Covaris S2 (Covaris, Woburn, MA, USA) for 8 min with the following set up: duty cycle: max, intensity: max, cycles/burst: max, mode: Power Tracking. The sonicated chromatin was diluted to 0.15% SDS and incubated overnight at 4°C with 10 μg of anti-BCL9 (Abcam, ab37305) or anti-TBX3m (Santacruz, sc-17871) or IgG and 50 ul of protein A/G magnetic beads (Upstate). The beads were washed at 4°C with wash buffer 1 (0.1% SDS, 0.1% deoxycholate, 1% Triton X-100, 0.15 M NaCl, 1 mM EDTA, 0.5 mM EGTA, 20 mM HEPES), wash buffer 2 (0.1% SDS, 0.1% sodium deoxycholate, 1% Triton X-100, 0.5 M NaCl, 1 mM EDTA, 0.5 mM EGTA, 20 mM HEPES), wash buffer 3 (0.25 M LiCl, 0.5% sodium deoxycholate, 0.5% NP-40, 1 mM EDTA, 0.5 mM EGTA, 20 mM HEPES), and finally twice with Tris EDTA buffer. The chromatin was eluted with 1% SDS, 0.1 M NaHCO3, de-crosslinked by incubation at 65°C for 5 hr with 200 mM NaCl, extracted with phenol-chloroform, and ethanol precipitated. The immunoprecipitated DNA was used as input material for DNA deep sequencing. The pull downs of BCL9 and TBX3 were performed in parallel experiments. Note that the ChIP-seq of BCL9 has been already described in Salazar et al., 2019, and entirely re-analyzed here (see below).

#### Data analysis

Overall sequencing quality of the acquired fastq files was assessed using FastQC (version 0.11.5). Because all the samples exhibited good quality (MAPQ > 30) and had no adapter contamination > 0.1% trimming of reads was not deemed necessary. In addition, test alignments against several reference genomes were done using the FastQ Screen tool (version 0.13.0). Reference genomes were downloaded from UCSC (http://hgdownload.cse.ucsc.edu/goldenpath/mm10/bigZips/). Quality results were summarized using MultiQC (version 1.7). Fastq files were mapped to mouse reference genome (mm10) using the read aligner Bowtie2 (version 2.3.4.1). The resulting alignment files were then adjusted (conversion to binary format, removal of read aligned to mitochondrial DNA and indexing) using SamTools (version 1.9). To identify genomic regions enriched with aligned reads the peak calling tool MACS2 (version 2.2.6) was used. Calculated p-values were adjusted for false discovery rate (FDR) using Benjamini-Hochberg procedure, generating q-values. A cutoff q < 0.05 was used to assess significance. IgG sample were used as enrichment-normalization control. MACS2 generated peak files were further filtered by removal of blacklisted regions according to the ENCODE project using bedtools (version 2.26.0). Annotation and visualization were made with the R programming language (version 3.4.4) and Rstudio (version 1.1.463), using R packages ChIPpeakAnno (version 3.12.7), ChIPseeker (version 1.14.2) and ggplot2 (version 3.2.1). CIRCOS plots was produced using the R package Circlize (version 0.4.8) and genomic track visualization was done with Integrative Genomics Viewer (IGV) (version 2.4.17). Motif analysis was performed with HOMER. The data have been deposited at ArrayExpress with accession number E-MTAB-8997.

### RNA-seq data analysis

Quality control of fastq files were done using FastQC (version 0.11.5). Trimming of reads to remove adapter remnants and low quality read (MAPQ < 30) was performed with BBDuk, part of the BBMap suite (version 38.58). Test alignments against different reference genomes were done with the FastQ Screen tool (version 0.13.0).

Reference genomes were downloaded from UCSC (http://hgdownload.cse.ucsc.edu/goldenpath/mm10/bigZips/). Quality results were summarized using MultiQC (version 1.7). Sequenced reads were mapped to mouse reference genome (mm10) using the Spliced Transcript Alignment to a Reference (STAR) software (version 2.7.3a). Reference genome data (FASTA and GTF) were downloaded from GENCODE, release M24 (https://www.gencodegenes.org/mouse/). Annotation and visualization were made with the R programming language (version 3.4.4) and Rstudio (version 1.1.463). Hierarchical clustered heatmap was produced with the pheatmap R package, using Ward’s Hierarchical Agglomerative Clustering Method. A complete list of bioinformatic tools and references is listed in the accompanying Supplementary Table. The RNA-seq experiment has been deposited at ArrayExpress with accession number E-MTAB-9000.

### Protein Immunoprecipitation and Mass Spectrometry

Dissected mouse tumors were minced in cold PBS and treated with a hypotonic lysis buffer (20 mM tris-HCl, 75 mM NaCl, 1.5 mM MgCl2, 1 mM EGTA, 0.5% NP-40, and 5% glycerol). Protein extracts obtained were incubated with 1 μg of anti-BCL9 antibody (ab54833 or ab37305, Abcam) and protein A–conjugated Sepharose beads (GE Healthcare); they were then diluted in lysis buffer to a final volume of 1 ml. After 4 hours of incubation at 4°C on a rotating wheel, the beads were spun down and washed three times in lysis buffer. All steps were performed on ice, and all buffers were supplemented with fresh protease inhibitors (cOmplete, Roche) and 1 mM phenylmethylsulfonyl fluoride. For detecting the proteins in Western blot, the pulldown reactions were treated with Laemmli buffer, boiled at 95°C for 15 min, and subjected to SDS-PAGE separation and blotting on a polyvinylidene difluoride (PVDF) membrane. The PVDF membrane was probed with the anti–TBX3.

For liquid chromatography–MS/MS analysis, the protein samples, already dissolved in Laemmli buffer, were submitted to a filter-aided sample preparation (FASP) and digested with trypsin in 100 mM triethylammonium bicarbonate buffer overnight. Desalted samples were dried completely in a vacuum centrifuge and reconstituted with 50 μl of 3% acetonitrile and 0.1% formic acid. Each peptide solution (4 μl) was analyzed on both Q Exactive and Fusion mass spectrometers (Thermo Scientific) coupled to EASY-nLC 1000 (Thermo Scientific). Spectra acquisition and peptide count was performed as described in details in Cantù et al., 2017.

### Cell culture

Human cancer cell lines (HEK293T -parental, ΔCTNNB1 and ΔTCF/LEF (Doumpas et al., 2018) - and HCT116) were cultured in DMEM (Thermo Fisher Scientific, Belmont, Massachusetts, US) supplemented with 10% fetal bovine serum (Gibco, Gaithersburg, USA) 1% L-glutamine and 1% penicillin-streptomycin at 37°C in a humidified chamber supplemented with 5% CO2. CHIR 99021 compound (Tocris Bioscience) was used as Wnt signalling activator. At 6 hours after transfection, the transfected cells were treated for 16 hours either with DMSO (0 µM Chir), 1 µM or 5 uM Chir. Following that, cells were collected for further experiments.

### Plasmids and transfection

TBX3-3xflag (donated by Peter J. Hurlin), pCMV-Pygo2-HA (Cantù et al., 2013), pCDNA3.1-BCL9-GFP (kindly gifted by Mariann Bienz), pcDNA3-eGFP (kindly provided by Stefan Koch), M50 SUPER 8X TOPFLASH pTA-Luc (Addgene 12456), M51 SUPER 8X FOPFLASH pTA-Luc (Addgene 12456) were transfected using Lipofectamine 2000 (Invitrogen, USA) following the manufacturer’s instructions. Twenty-four hours after transfection, the cells were collected for the indicated experiments.

### RNA extraction and quantitative reverse transcriptase PCR (qRT-PCR)

Total RNAs from cells were extracted using TRIzol (Thermo Fisher Scientific, Belmont, Massachusetts, US) following the manufacturer’s instructions. After reverse transcription reaction with High Capacity cDNA Reverse Transcription Kit (Thermo Fisher Scientific), qPCR was conducted with SYBR green LightCycler 480 (Roche) on CFX96™ Real-Time PCR Detection System (Bio-Rad, USA). *GAPDH* was used as endogenous control and the relative expression of RNAs was calculated using the 2−ΔΔCt method. The primers used in this study were designed by Primer3plus web and their sequences are available upon request.

### Xenograft tumor zebrafish model

HCT116 cells were transfected with pCS2 (empty vector) or TBX3-expressing plasmid and labeled with 1,1′-dioctadecyl-3,3,3′,3′-tetramethylindocarbocyanine perchlorate (DiI, cat. no. D3899; Invitrogen) according to previously described method (Rouhi et al., 2010). Briefly, tumor cells were washed twice with PBS, followed by labeling with a final concentration of 5 μg/ml of DiI for 30 min at 37 °C. After rigorous washing with PBS, tumor cells were trypsinized for 5 minutes, counted under a phase contrast microscope, centrifuged at 300 g for 5 min and resuspended at a final concentration of 30–40 cells per µl in medium. Human cells were injected into the perivitelline space of 72 h old fli1:EGFP transgenic zebrafish embryos. The eggs were fertilized, collected and dechorionated. After that, HCT116 cells were injected into the vitreous cavity of zebrafish embryo with non-filamentous borosilicate glass capillary needles attached to the microinjector under the stereomicroscope (Leica Microsystems). After tumor cell injection, zebrafish embryos were further selected under fluorescent microscopy to ensure that tumor cells were located only within the cavity and then incubated in aquarium water for consecutive 3 days at 36.0 °C. Primary tumor growth, invasion and metastasis in the zebrafish body were monitored at day 3 with a fluorescent microscope (Nikon Eclipse C1) as previously published (Rouhi et al., 2010). Briefly, each zebrafish embryo was picked up and monitored to detect tumor cell distribution. Two different sets of images from the head region and the trunk region were collected separately from each zebrafish embryo. Disseminated tumor cells in the caudal hematopoietic plexus of zebrafish embryos were counted (in a double-blind manner) and the primary tumor areas were measured using Image J software. At least 15–20 embryos were included in each experimental group.

### Western blot

Cells were lysed at 4°C for 10 min using NP-40 lysis buffer (50 mM Tris-HCl (pH 7.5), 120 mM NaCl, and 1% (v/v) Igepal), supplemented with protease inhibitor cocktail (Merck). Cells were sonicated with Q700CA Sonicator (Q Sonica) on 50 amplitude for 5 sec on/off for 2 cycles. Protein lysates were separated by 10% SDS-polyacrylamide gel electrophoresis (SDS-PAGE) and transferred onto nitrocellulose membrane. The membrane was blocked in Odyssey TBS Blocking Buffers (LI-COR Bioscience) for 1 h at room temperature and subsequently incubated with specific antibody against FLAG (1:1000, Merck F3165) overnight at 4°C and HA (1:1000, Merck 05-902R) and GFP (1:1000 Santa Cruz Biotechnology sc-9996) 1 h at room temperature. Afterwards, the membrane was washed and incubated with fluorescent Goat-anti-Rabbit and goat-anti-Mouse (1:2000; LI-COR Bioscience) for 1 h at room temperature. Protein detection was performed in LICOR Odyssey CLx (LI-COR Bioscience).

### Super-TOP/FOP-Flash reporter assay

The β-catenin reporter plasmid (TOP-flash) and its mutant control (FOP-flash) were constructed by Addgene (12456). Cells were plated in triplicate in a 96 well plated overnight and co-transfected with 50 ng TOP flash or FOP flash expression plasmids together with other plasmids for the experiments using Lipofectamine 2000. The activities of firefly luciferase reporters were determined at 24 hours after transfection using ONE-Step Luciferase Assay System Kit (Pierce) according to the manufacturer’s instructions. The TOP/FOP ratio was used as a measure of β-catenin-driven transcription. GraphPad Prism software was used for statistical analyses. Values were expressed as mean ± standard deviation. Mann Whitney U test was used to analyze the differences between two groups and *P* < 0.05 was considered statistically significant. All experiments were performed at least three times and all samples analyzed in triplicate unless otherwise stated.

## Acknowledgements

The authors thank Peter J. Hurlin for kindly donating the *TBX3* expressing plasmid, George Hausmann, Stefan Koch, Lavanya Moparthi for sharing reagents and critical input, and Lasse Jensen for helping with zebrafish injection. C.C. is a Wallenberg Molecular Medicine fellow and receives generous financial support from the Knut and Alice Wallenberg Foundation. This work was supported by Cancerfonden to C.C. (CAN 2018/542), by the Swiss National Science Foundation and the Canton of Zurich to K.B. and by the Swiss National Science Foundation (grant PCEPP3_181249) to AEM.

## Author contribution

DZ, CB, AJM, FMS, AEM and CC performed the experiments and designed the figures. SS analyzed the RNA-seq and ChIP-seq data and designed the respective figure panels. MA, SB, ND and JR assisted with the experiments or their design. KB, AEM and CC conceived the project, obtained the funding and supervised the research teams. AEM and CC wrote the manuscript in Malta.

---

**Figure supplement 1:**
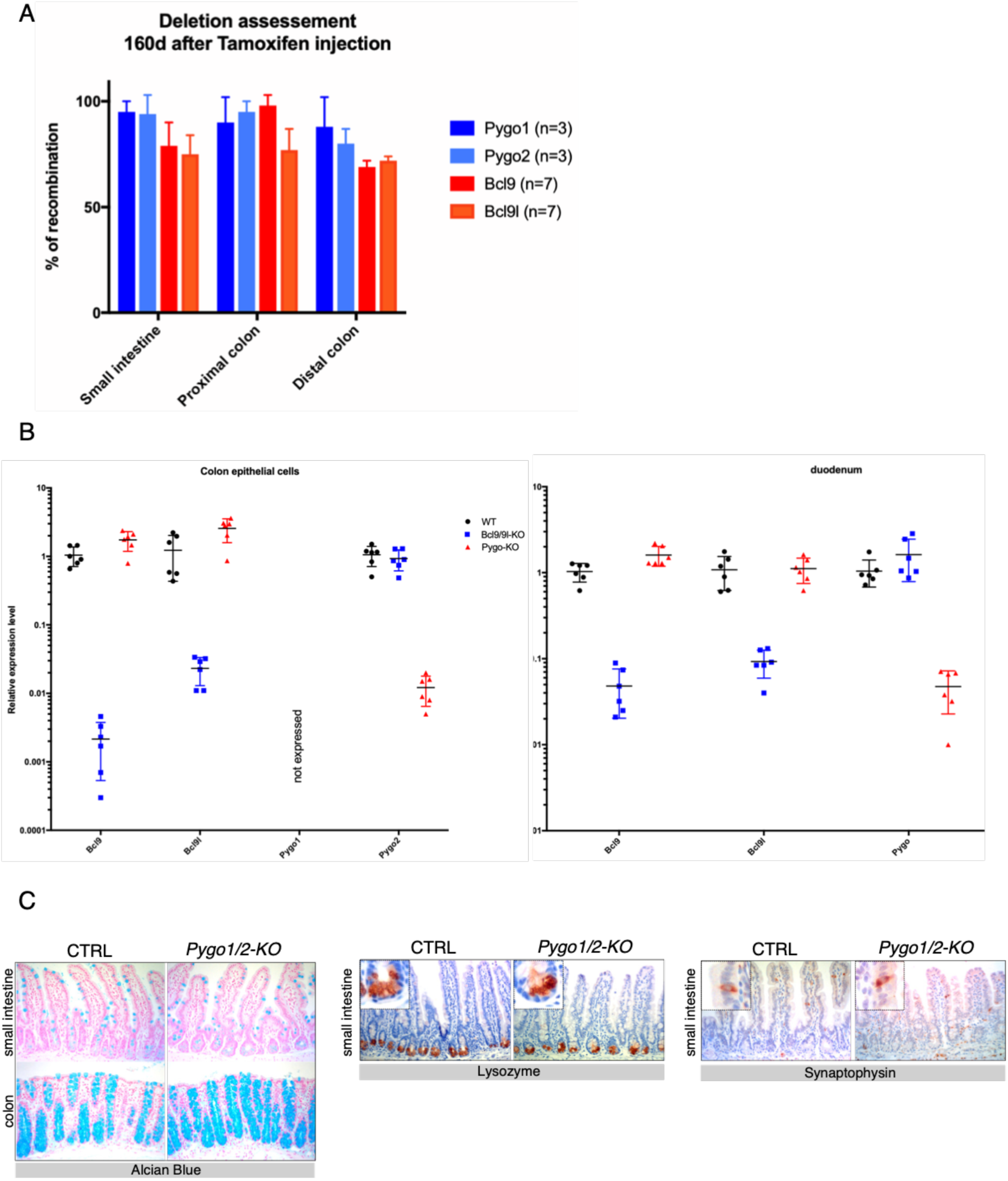
(**A**) Specific deletion of *Bcl9/9*l and *Pygo1/2* in the intestinal epithelium was achieved by introducing a tamoxifen-dependent vil-Cre-ERt2 driver. Mice were injected with tamoxifen at four weeks of age and analyzed at different time-points thereafter. Deletion was monitored by real-time PCR on genomic DNA. Gene deletion was stable over at least 23 weeks, indicating that the stem cell compartment was successfully hit, and that no selective disadvantage of mutant compared to wild-type cells occurred. (**B**) *Bcl9/9l* and *Pygo1/2* deletion was also monitored via qRT-PCR both in colon (left panel) and in duodenal cells (right panel). (**C**) Histological analysis was performed on the small intestine and the colon. Compared to control (CTRL) littermates, conditional *Pygo1/2-*KO mutants displayed normal intestinal architecture. We detected all differentiated cell types, including the secretory lineages such as goblet cells (alcian blue positive), paneth cells (lysozyme positive), and enteroendocrine cells (synaptophysin positive).

**Figure supplement 2:**
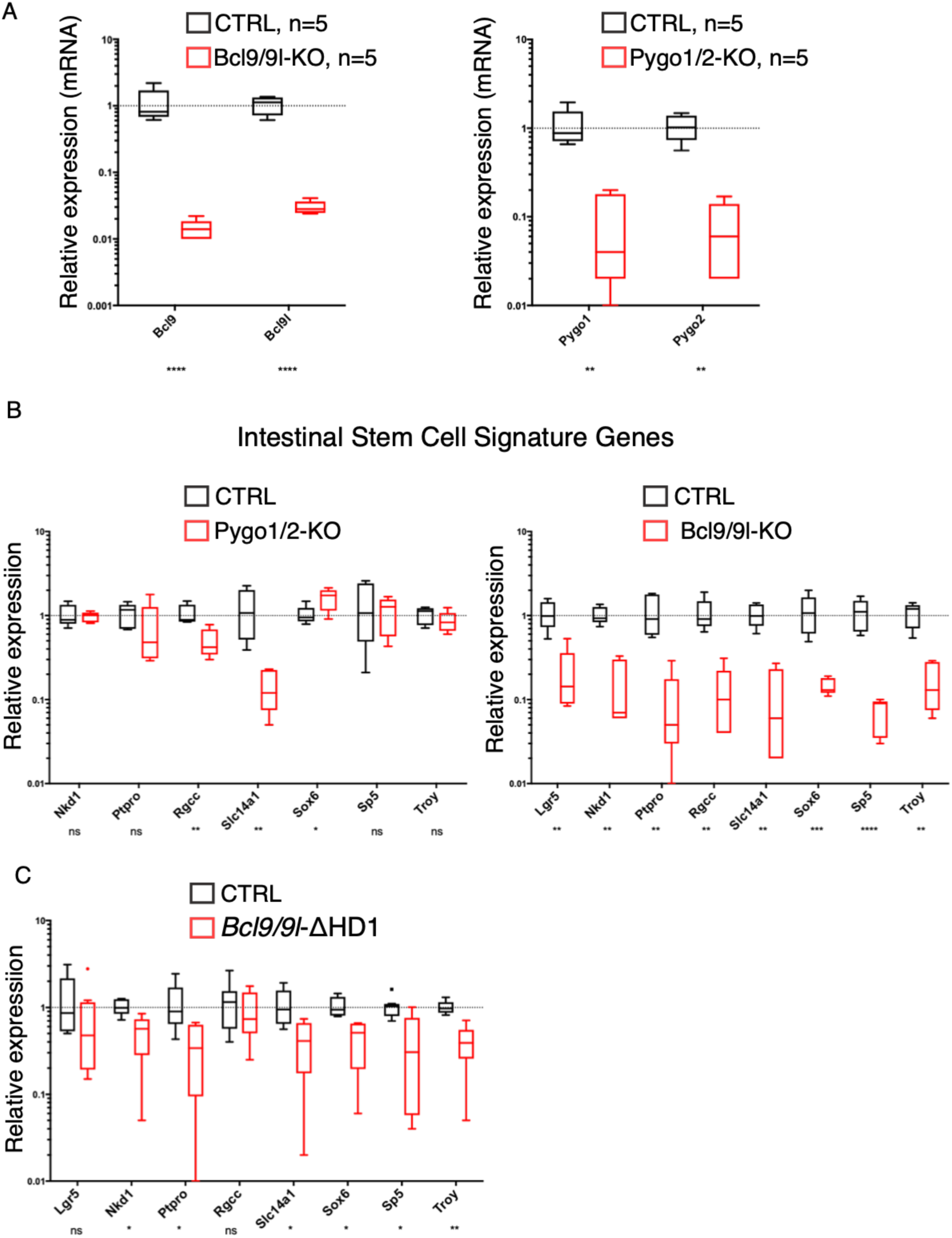
(**A**) Genetic recombination of *Bcl9/9l* and *Pygo1/2* in tumors was monitored via qRT-PCR. Their expression is dramatically reduced (red bars) when compared to that in control mice (CTRL, black bars) confirming high recombination rate. (**B**) The intestinal epithelium-specific recombination of *Pygo1/2* (in *Pygo1/2*-KO) does not recapitulate the effects of deleting *Bcl9/9l* (*Bcl9/9l*-KO) in dysplastic adenomas. The expression of genes associated with intestinal stem cell function was analyzed via quantitative RT-PCR, and the expression in the mutant cells (red) was compared to that in the control (CTRL, black). (**C**) Quantitative RT-PCR of target genes associated with intestinal stem cell function (compare it with the same analysis of *Pygo1/2-*KO in panel B) of RNA extracted from control (CTRL, black) or Bcl9/9l-ΔHD1 (red) tumors.

**Figure supplement 3:**
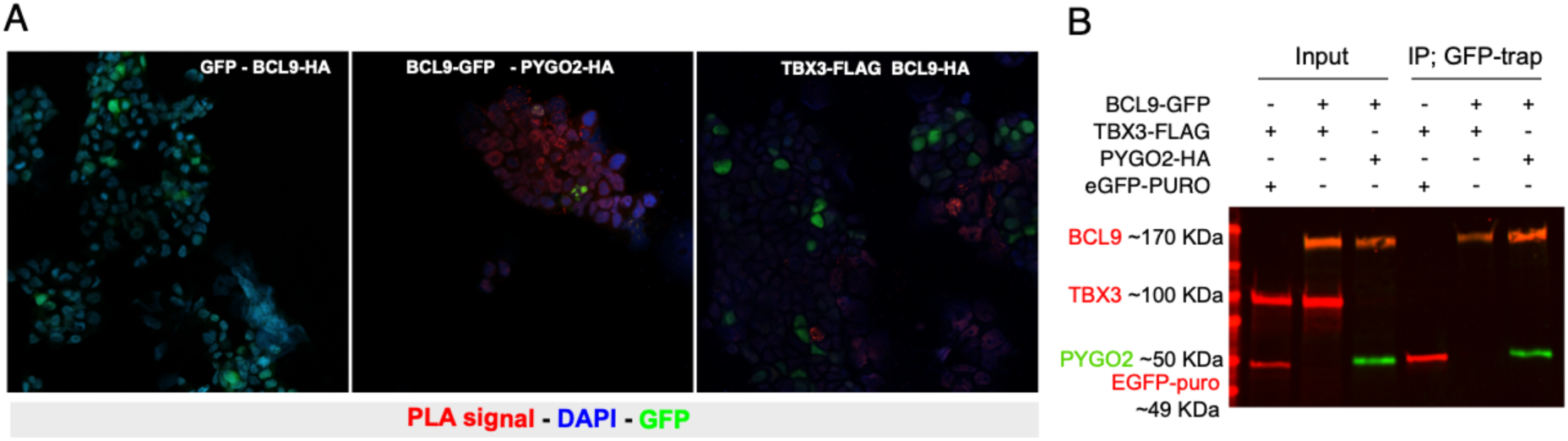
(**A**) Proximity Ligation Assay (PLA) confirms vicinity between BCL9 and TBX3 when expressed in human HEK293T cells: red signal is observed upon protein-protein proximity. Nuclear GFP and BCL9 do not give rise to PLA signal (left panel, negative control). PYGO2 is used as positive interactor with BCL9 (middle panel) and gives rise to PLA signal (red). TBX3 and BCL9 produce nuclear PLA signal in a subset of transfected cells (detected with GFP; right panel). (**B**) Immunoprecipitation assay using a GFP-trap strategy. BCL9 reliably interacts with HA-tagged PYGO2 in high-salt washing conditions (400 mM NaCl). In this stringent condition, FLAG-tagged TBX3 is not observed when GFP-BCL9 is immunoprecipitated, suggesting absence of direct interaction or affinity inferior to that between BCL9 and PYGO2. IP of GFP is used a control.

**Figure supplement 4:**
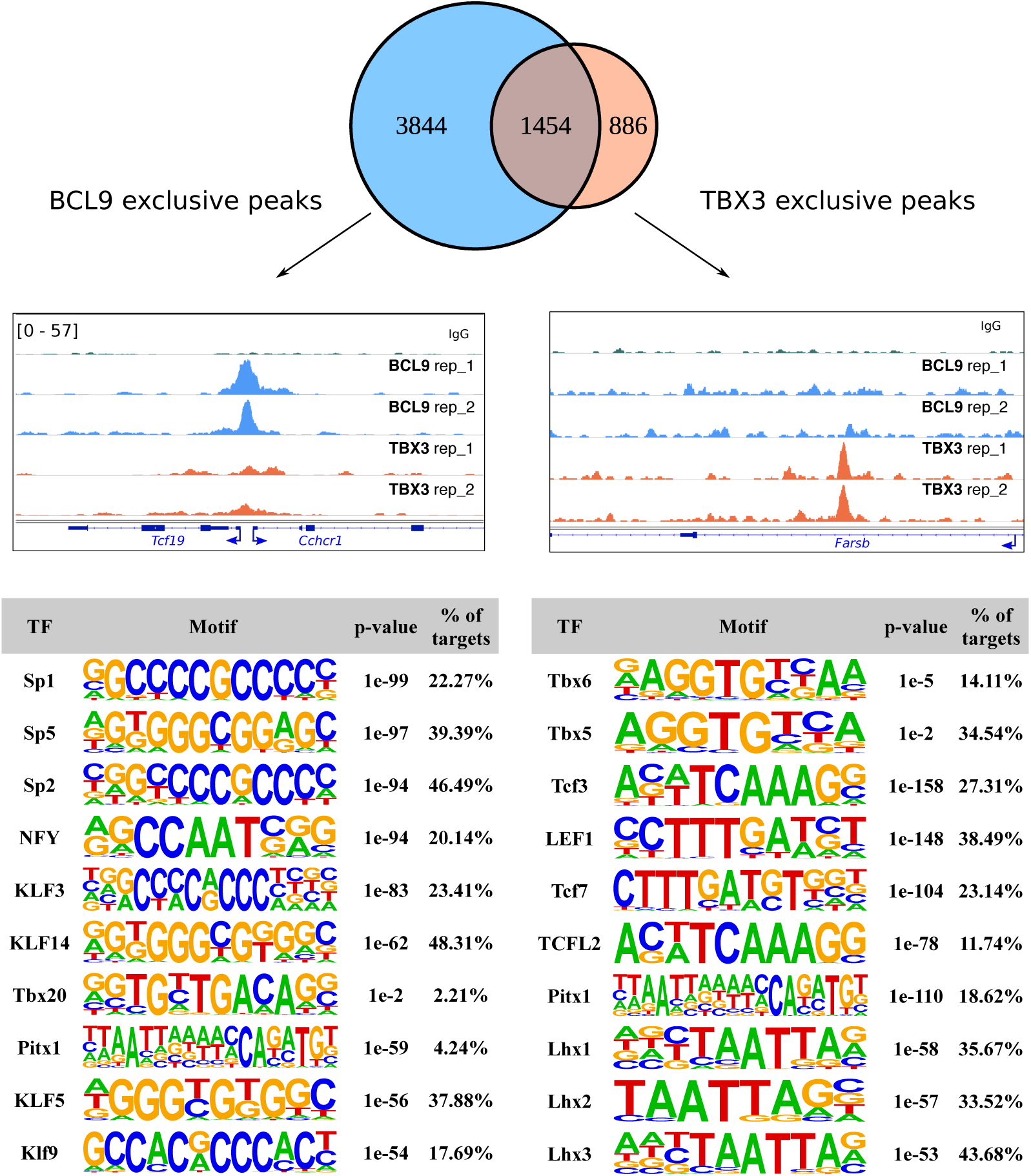
Overlap of the high-confidence BCL9 and TBX3 peaks in developing murine limbs reveals the existence of BCL9 exclusive (one example displayed in the genomic tracks on the left) and TBX3 exclusive peaks (one example in the genomic track on the right). The greatest group of BCL9 exclusive peaks, surprisingly, does not display high enrichment of TCF/LEF motifs, suggesting that a predominant genomic function of BCL9 is not related to Wnt/β-catenin signalling, as previously suggested by our work and that of others (Cantù et al., 2018, 2017, 2014; Jiang et al., 2020). On the other hand, BCL9 exclusive peaks (bottom-left table) show massive enrichment of Zinc Finger containing Sp/Krüppel-like-Factor (KLF) family of transcription factors. The TBX3 exclusive peaks (bottom-right table) display presence of TBX motifs – absent in the BCL9/TBX3 overlapping peaks. Genomic tracks are adapted for this figure from IGV Integrative Genomic Viewer (https://igv.org/). Motif analysis was performed with HOMER (http://homer.ucsd.edu/homer/).

**Figure supplement 5:**
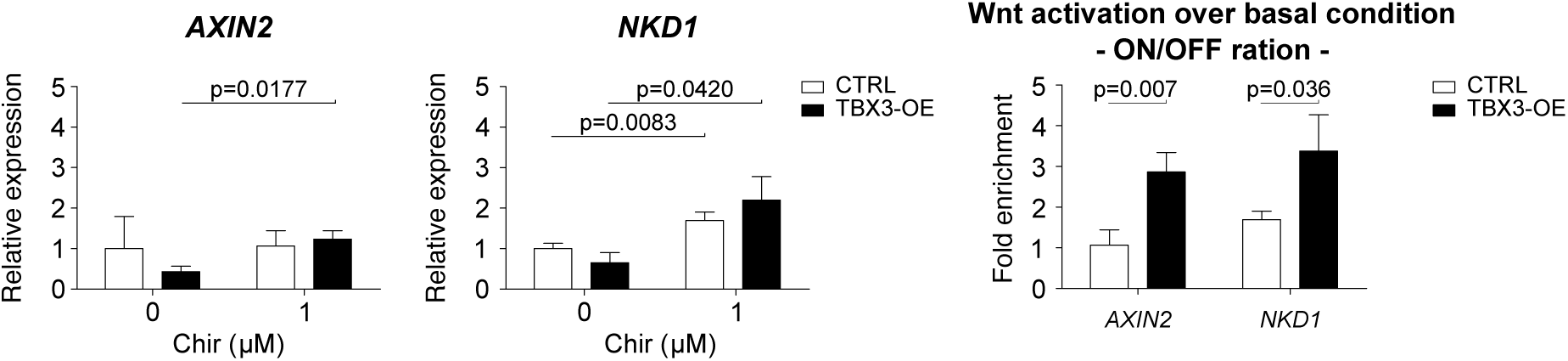
*AXIN2* and *NKD1* are here considered as representative Wnt transcriptional targets. mRNA expression in HEK293T cells upon TBX3 overexpression (black bars) is measured via qRT-PCR and compared to control condition (CTRL, transfection with empty vector, white bars). Consistently with the STF/Luciferase assay (Figure 4C), TBX3 represses the transcription of endogenous Wnt targets. Upon suboptimal Wnt stimulation however (achieved via addition of 1 μM of CHIR), *AXIN2* and *NKD1* are only mildly activated in CTRL condition. On the other hand, TBX3 overexpression induces a substantial statistical increase in *AXIN2* and *NKD1* mRNA abundance; the Wnt-ON (CHIR 1 µM) condition is compared to the Wnt-OFF basal condition (ON/OFF ration; right panel). The data are presented as the mean ± SD of three independent experiments. Parametric unpaired Student t-test was used to measure the statistical significance (p-values above the charts). *GAPDH* mRNA was used to normalize transcript levels.

## References

Barker N, van Es JH, Kuipers J, Kujala P, van den Born M, Cozijnsen M, Haegebarth A, Korving J, Begthel H, Peters PJ, Clevers H. 2007. Identification of stem cells in small intestine and colon by marker gene Lgr5. Nature 449:1003–7. doi:10.1038/nature06196

Cantù C, Felker A, Zimmerli D, Prummel KD, Cabello EM, Chiavacci E, Méndez-Acevedo KM, Kirchgeorg L, Burger S, Ripoll J, Valenta T, Hausmann G, Vilain N, Aguet M, Burger A, Panáková D, Basler K, Mosimann C. 2018. Mutations in Bcl9 and Pygo genes cause congenital heart defects by tissue-specific perturbation of Wnt/β-catenin signaling. Genes Dev 32:1443–1458. doi:10.1101/gad.315531.118

Cantù C, Pagella P, Shajiei TD, Zimmerli D, Valenta T, Hausmann G, Basler K, Mitsiadis TA. 2017. A cytoplasmic role of Wnt/β-catenin transcriptional cofactors Bcl9, Bcl9l, and Pygopus in tooth enamel formation. Sci Signal 10:1–11. doi:10.1126/scisignal.aah4598

Cantù C, Valenta T, Hausmann G, Vilain N, Aguet M, Basler K. 2013. The Pygo2-H3K4me2/3 interaction is dispensable for mouse development and Wnt signaling-dependent transcription. Development 140:2377–86. doi:10.1242/dev.093591

Cantù C, Zimmerli D, Hausmann G, Valenta T, Moor A, Aguet M, Basler K. 2014. Pax6-dependent, but beta-catenin-independent, function of Bcl9 proteins in mouse lens development. Genes Dev 1879–1884. doi:10.1101/gad.246140.114

Capdevila J, Tsukui T, Esteban CR, Zappavigna V, Belmonte JCI. 1999. Control of vertebrate limb outgrowth by the proximal factor Meis2 and distal antagonism of BMPs by Gremlin. Mol Cell 4:839–849. doi:10.1016/S1097-2765(00)80393-7

Deka J, Wiedemann N, Anderle P, Murphy-Seiler F, Bultinck J, Eyckerman S, Stehle J-C, André S, Vilain N, Zilian O, Robine S, Delorenzi M, Basler K, Aguet M. 2010. Bcl9/Bcl9l are critical for Wnt-mediated regulation of stem cell traits in colon epithelium and adenocarcinomas. Cancer Res 70:6619–28. doi:10.1158/0008-5472.CAN-10-0148

Doumpas N, Lampart F, Robinson MD, Lentini A, Nestor CE, Cantù C, Basler K. 2018. TCF/LEF dependent and independent transcriptional regulation of Wnt/β-catenin target genes. EMBO J 38:e98873. doi:10.15252/embj.201798873

Fiedler M, Graeb M, Mieszczanek J, Rutherford TJ, Johnson CM, Bienz M. 2015. An ancient Pygo-dependent Wnt enhanceosome integrated by chip/LDB-SSDP. Elife 4:1–22. doi:10.7554/eLife.09073

Frank DU, Emechebe U, Thomas KR, Moon AM. 2013. Mouse Tbx3 Mutants Suggest Novel Molecular Mechanisms for Ulnar-Mammary Syndrome. PLoS One 8:1–7. doi:10.1371/journal.pone.0067841

Gay DM, Ridgway RA, Müeller M, Hodder MC, Hedley A, Clark W, Leach JD, Jackstadt R, Nixon C, Huels DJ, Campbell AD, Bird TG, Sansom OJ. 2019. Loss of BCL9/9l suppresses Wnt driven tumourigenesis in models that recapitulate human cancer. Nat Commun 10:1–16. doi:10.1038/s41467-019-08586-3

Grifone R, Demignon J, Giordani J, Niro C, Souil E, Bertin F, Laclef C, Xu PX, Maire P. 2007. Eya1 and Eya2 proteins are required for hypaxial somitic myogenesis in the mouse embryo. Dev Biol 302:602–616. doi:10.1016/j.ydbio.2006.08.059

Jiang M, Kang Y, Sewastianik T, Wang J, Tanton H, Alder K, Dennis P, Xin Y, Wang Z, Liu R, Zhang M, Huang Y, Loda M, Srivastava A, Chen R, Liu M, Carrasco RD. 2020. BCL9 provides multi-cellular communication properties in colorectal cancer by interacting with paraspeckle proteins. Nat Commun 11. doi:10.1038/s41467-019-13842-7

Kahn M. 2014. Can we safely target the WNT pathway? Nat Rev Drug Discov 13:513–532.

Kennedy MW, Cha S-W, Tadjuidje E, Andrews PG, Heasman J, Kao KR. 2010. A co-dependent requirement of xBcl9 and Pygopus for embryonic body axis development in Xenopus. Dev Dyn 239:271–83. doi:10.1002/dvdy.22133

Kim JJ, Shajib MS, Manocha MM, Khan WI. 2012. Investigating intestinal inflammation in DSS-induced model of IBD. J Vis Exp 1–6. doi:10.3791/3678

Kramps T, Peter O, Brunner E, Nellen D, Froesch B, Chatterjee S, Murone M, Züllig S, Basler K. 2002. Wnt/wingless signaling requires BCL9/legless-mediated recruitment of pygopus to the nuclear beta-catenin-TCF complex. Cell 109:47–60.

Li B, Rheaume C, Teng A, Bilanchone V, Munguia JE, Hu M, Jessen S, Piccolo S, Waterman ML, Dai X, Rhe C, Teng A, Bilanchone V, Munguia JE, Hu M, Jessen S, Piccolo S, Waterman ML, Dai X. 2007. Developmental phenotypes and reduced Wnt signaling in mice deficient for pygopus 2. Genesis 45:318–325. doi:10.1002/dvg

Li D, Sakuma R, Vakili NA, Mo R, Puviindran V, Deimling S, Zhang X, Hopyan S, Hui C chung. 2014. Formation of proximal and anterior limb skeleton requires early function of Irx3 and Irx5 and is negatively regulated by shh signaling. Dev Cell 29:233–240. doi:10.1016/j.devcel.2014.03.001

Lyou Y, Habowski AN, Chen GT, Waterman ML. 2017. Inhibition of nuclear Wnt signalling: challenges of an elusive target for cancer therapy. Br J Pharmacol 174:4589–4599. doi:10.1111/bph.13963

Mani M, Carrasco DE, Zhang Y, Takada K, Gatt ME, Dutta-Simmons J, Ikeda H, Diaz-Griffero F, Pena-Cruz V, Bertagnolli M, Myeroff LL, Markowitz SD, Anderson KC, Carrasco DR. 2009. BCL9 promotes tumor progression by conferring enhanced proliferative, metastatic, and angiogenic properties to cancer cells. Cancer Res 69:7577–86. doi:10.1158/0008-5472.CAN-09-0773

Mieszczanek J, van Tienen LM, Ibrahim AEK, Winton DJ, Bienz M. 2019. Bcl9 and Pygo synergise downstream of Apc to effect intestinal neoplasia in FAP mouse models. Nat Commun 10. doi:10.1038/s41467-018-08164-z

Moor AE, Anderle P, Cantù C, Rodriguez P, Wiedemann N, Baruthio F, Deka J, André S, Valenta T, Moor MB, Győrffy B, Barras D, Delorenzi M, Basler K, Aguet M. 2015. BCL9/9L-β-catenin Signaling is Associated With Poor Outcome in Colorectal Cancer. EBioMedicine. doi:10.1016/j.ebiom.2015.10.030

Mosimann C, Hausmann G, Basler K. 2009. Beta-catenin hits chromatin: regulation of Wnt target gene activation. Nat Rev Mol Cell Biol 10:276–86. doi:10.1038/nrm2654

Mouradov D, Sloggett C, Jorissen RN, Love CG, Li S, Burgess AW, Arango D, Strausberg RL, Buchanan D, Wormald S, O’Connor L, Wilding JL, Bicknell D, Tomlinson IPM, Bodmer WF, Mariadason JM, Sieber OM. 2014. Colorectal cancer cell lines are representative models of the main molecular subtypes of primary cancer. Cancer Res 74:3238–3247. doi:10.1158/0008-5472.CAN-14-0013

Nusse R, Clevers H. 2017. Wnt/β-Catenin Signaling, Disease, and Emerging Therapeutic Modalities. Cell 169:985–999. doi:10.1016/j.cell.2017.05.016

Parker DS, Jemison J, Cadigan KM. 2002. Pygopus, a nuclear PHD-finger protein required for Wingless signaling in Drosophila. Development 129:2565–76.

Rouhi P, Jensen LD, Cao Z, Hosaka K, Länne T, Wahlberg E, Steffensen JF, Cao Y. 2010. Hypoxia-induced metastasis model in embryonic zebrafish. Nat Protoc 5:1911–1918. doi:10.1038/nprot.2010.150

Salazar VS, Capelo LP, Cantù C, Zimmerli D, Gamer L, Ohte S, Economides A, Carey DJ, Feigenson M, Pregizer S, Nyman JS, Gosalia N, Cox K, Basler K, Rosen V. 2019. Reactivation of a developmental Bmp2 signaling center is required for therapeutic control of the periosteal niche in the murine skeleton. Elife 8:1–31. doi:10.7554/elife.42386

Schwab KR, Patterson LT, Hartman HA, Song N, Lang RA, Lin X, Potter SS. 2007. Pygo1 and Pygo2 roles in Wnt signaling in mammalian kidney development. BMC Biol 5:15. doi:10.1186/1741-7007-5-15

Talla SB, Brembeck FH. 2016. The role of *Pygo2* for Wnt/&#xDF;-catenin signaling activity during intestinal tumor initiation and progression. Oncotarget 7:80612–80632. doi:10.18632/oncotarget.13016

Valenta T, Hausmann G, Basler K. 2012. The many faces and functions of β-catenin. EMBO J 31:2714–36. doi:10.1038/emboj.2012.150

van Tienen LM, Mieszczanek J, Fiedler M, Rutherford TJ, Bienz M, Labhart T, Desplan C, Hursh D, Jones T, Bejsovec A, Peifer M, Mortin M, Clevers H, Tolstorukov M, Luquette L, Xi R, Jung Y, Park R, Bishop E, Canfield T, Sandstrom R, Thurman R, MacAlpine D, Stamatoyannopoulos J, Kellis M, Elgin S, Kuroda M, Pirrotta V, Karpen G, Park P. 2017. Constitutive scaffolding of multiple Wnt enhanceosome components by Legless/BCL9. Elife 6:477–488. doi:10.7554/eLife.20882

Willmer T, Cooper A, Peres J, Omar R, Prince S. 2017. The T-Box transcription factor 3 in development and cancer. Biosci Trends 11:254–266. doi:10.5582/bst.2017.01043

Zimmerli D, Hausmann G, Cantù C, Basler K. 2017. Pharmacological interventions in the Wnt pathway: Inhibition of Wnt secretion versus disrupting the protein-protein interfaces of nuclear factors. Br J Pharmacol. doi:10.1111/bph.13864

