## Supplementary Table for "TBX3 acts as tissue-specific component of the Wnt/β-catenin enhanceosome"

### Supplementary Table – Bioinformatics Resources

The following table present key resources used for the computational analysis of RNA-seq and ChIP-seq data. References to corresponding publications and links to online sources are included. Python-based software and tools have been obtained from the Anaconda Cloud (https://anaconda.com) using the Bioconda channel (https://bioconda.github.io/). R-based packages have been obtained from either the Comprehensive R Archive Network (CRAN) or from Bioconductor (https://bioconductor.org/about/).

| Resource/Software/Algorithm | References | Available |
|---|---|---|
| **Mouse reference genome (mm10), UCSC** | - | (http://hgdownload.cse.ucsc.edu/goldenpath/mm10/bigZips/). |
| **Mouse reference genome and annotation, release M24, GENCODE** | (Frankish et al., 2019) | https://www.gencodegenes.org/mouse/ |
| **ENCODE blacklisted regions, Version 2** | (Amemiya et al., 2019) | https://github.com/Boyle-Lab/Blacklist/tree/master/lists |
| **R programming language Version 3.4.4** | (R Core Team, 2017) | https://cran.r-project.org/ |
| **Rstudio Version 1.1.463** | (Rstudio Team, 2015) | https://rstudio.com/ |
| **FastQC Version 0.11.5** | (Andrews, 2010) | https://anaconda.org/bioconda/fastqc |
| **FastQ Screen Version 0.13.0** | (Wingett and Andrews, 2018) | https://anaconda.org/bioconda/fastq-screen |
| **MultiQC Version 1.7** | (Ewels et al., 2016) | https://anaconda.org/bioconda/multiqc |
| **Bowtie2 Version 2.3.4.1** | (Langmead and Salzberg, 2012) | https://anaconda.org/bioconda/bowtie2 |
| **SamTools Version 1.9** | (Li et al., 2009) | https://anaconda.org/bioconda/samtools |
| **MACS2 Version 2.2.6** | (Zhang et al., 2008) | https://anaconda.org/bioconda/macs2 |
| **BedTools Version 2.26.0** | (Quinlan and Hall, 2010) | https://anaconda.org/bioconda/bedtools |
| **ChIPpeakAnno, R-package Version 3.12.7** | (Zhu et al., 2010) | https://www.bioconductor.org/packages/3.6/bioc/html/ChIPpeakAnno.html |
| **ChIPseeker, R-package Version 1.14.2** | (Yu et al., 2015) | https://www.bioconductor.org/packages/3.6/bioc/html/ChIPseeker.html |
| **ggplot2, R-package Version 3.2.1** | (Wickham, 2016) | https://ggplot2.tidyverse.org/ |

| | | |
|---|---|---|
| **Circlize, R-package Version 0.4.8** | (Gu et al., 2014) | https://cran.r-project.org/web/packages/circlize/index.html |
| **Integrative Genomic Viewer (IGV), Version 2.4.17** | (Robinson et al., 2011) | https://anaconda.org/bioconda/igv |
| **HOMER** | (Heinz et al., 2010) | https://anaconda.org/bioconda/homer |
| **BBDuk, part of the BBMap suite, Version 38.58** | (Bushnell, n.d.) | sourceforge.net/projects/bbmap |
| **Spliced Transcripts Alignment to a Reference (STAR), Version 2.7.3a** | (Dobin et al., 2013) | https://anaconda.org/bioconda/star |
| **Benjamini-Hochberg FDR correction (MACS2)** | (Benjamini and Hochberg, 2018) | - |
| **Pheatmap, R-package Version 1.0.12** | (Kolde, 2019) | https://CRAN.R-project.org/package=pheatmap |
| **GeneOverlap, R-package Version 1.14.0** | (Shen and Sinai, 2013) | http://shenlab-sinai.github.io/shenlab-sinai/ |

### Supplementary References


Amemiya HM, Kundaje A, Boyle AP. 2019. The ENCODE Blacklist: Identification of Problematic Regions of the Genome. *Sci Rep* **9**:9354. doi:10.1038/s41598-019-45839-z

Andrews S. 2010. FastQC: a quality control tool for high throughput sequence data. *Babraham Bioinforma*. doi:citeulike-article-id:11583827

Benjamini Y, Hochberg Y. 2018. Controlling the False Discovery Rate: A Practical and Powerful Approach to Multiple Testing. *J R Stat Soc Ser B* **57**:289–300. doi:10.1111/j.2517-6161.1995.tb02031.x

Bushnell B. n.d. BBDuk. *sourceforge.net*. sourceforge.net/projects/bbmap/

Dobin A, Davis CA, Schlesinger F, Drenkow J, Zaleski C, Jha S, Batut P, Chaisson M, Gingeras TR. 2013. STAR: Ultrafast universal RNA-seq aligner. *Bioinformatics* **29**:15–21. doi:10.1093/bioinformatics/bts635

Ewels P, Magnusson M, Lundin S, Käller M. 2016. MultiQC: summarize analysis results for multiple tools and samples in a single report. *Bioinformatics* **32**:3047–3048. doi:10.1093/bioinformatics/btw354

Frankish A, Diekhans M, Ferreira A-M, Johnson R, Jungreis I, Loveland J, Mudge JM, Sisu C, Wright J, Armstrong J, Barnes I, Berry A, Bignell A, Carbonell Sala S, Chrast J, Cunningham F, Di Domenico T, Donaldson S, Fiddes IT, García Girón C, Gonzalez JM, Grego T, Hardy M, Hourlier T, Hunt T, Izuogu OG, Lagarde J, Martin FJ, Martínez L, Mohanan S, Muir P, Navarro FCP, Parker A, Pei B, Pozo F, Ruffier M, Schmitt BM, Stapleton E, Suner M-M, Sycheva I, Uszczynska-Ratajczak B, Xu J, Yates A, Zerbino D, Zhang Y, Aken B, Choudhary JS, Gerstein M, Guigó R, Hubbard TJP, Kellis M, Paten B, Reymond A, Tress ML, Flicek P. 2019. GENCODE reference annotation for the human and mouse genomes. *Nucleic Acids Res* **47**:D766–D773. doi:10.1093/nar/gky955

Gu Z, Gu L, Eils R, Schlesner M, Brors B. 2014. circlize implements and enhances circular visualization in R. *Bioinformatics* **30**:2811–2812. doi:10.1093/bioinformatics/btu393

Heinz S, Benner C, Spann N, Bertolino E, Lin YC, Laslo P, Cheng JX, Murre C, Singh H, Glass CK. 2010. Simple Combinations of Lineage-Determining Transcription Factors Prime cis-Regulatory Elements Required for Macrophage and B Cell Identities. *Mol Cell* **38**:576–589. doi:10.1016/j.molcel.2010.05.004

Kolde R. 2019. pheatmap: Pretty Heatmaps. *R Packag version 1012*. https://cran.r-project.org/package=pheatmap

Langmead B, Salzberg SL. 2012. Fast gapped-read alignment with Bowtie 2. *Nat Methods* **9**:357–359. doi:10.1038/nmeth.1923

Li H, Handsaker B, Wysoker A, Fennell T, Ruan J, Homer N, Marth G, Abecasis G, Durbin R. 2009. The Sequence Alignment/Map format and SAMtools. *Bioinformatics* **25**:2078–2079. doi:10.1093/bioinformatics/btp352

Quinlan AR, Hall IM. 2010. BEDTools: A flexible suite of utilities for comparing genomic features. *Bioinformatics* **26**:841–842. doi:10.1093/bioinformatics/btq033

R Core Team. 2017. R: A language and environment for statistical computing.

Robinson JT, Thorvaldsdóttir H, Winckler W, Guttman M, Lander ES, Getz G, Mesirov JP. 2011. Integrative genomics viewer. *Nat Biotechnol* **29**:24–26. doi:10.1038/nbt.1754

Rstudio Team. 2015. RStudio: Integrated Development for R.

Shen L, Sinai M. 2013. GeneOverlap: Test and visualize gene overlaps. *R Packag version 1140*. http://shenlab-sinai.github.io/shenlab-sinai/

Wickham H. 2016. ggplot2: Elegant Graphics for Data Analysis. New York: Springer-Verlag.

Wingett SW, Andrews S. 2018. FastQ Screen: A tool for multi-genome mapping and quality control. *F1000Research* **7**:1338. doi:10.12688/f1000research.15931.2

Yu G, Wang L-G, He Q-Y. 2015. ChIPseeker: an R/Bioconductor package for ChIP peak annotation, comparison and visualization. *Bioinformatics* **31**:2382–2383. doi:10.1093/bioinformatics/btv145

Zhang Y, Liu T, Meyer CA, Eeckhoute J, Johnson DS, Bernstein BE, Nussbaum C, Myers RM, Brown M, Li W, Shirley XS. 2008. Model-based analysis of ChIP-Seq (MACS). *Genome Biol* **9**. doi:10.1186/gb-2008-9-9-r137

Zhu LJ, Gazin C, Lawson ND, Pagès H, Lin SM, Lapointe DS, Green MR. 2010. ChIPpeakAnno: A Bioconductor package to annotate ChIP-seq and ChIP-chip data. *BMC Bioinformatics* **11**:237. doi:10.1186/1471-2105-11-237
